## Supplemental Information for "A *Vaccinia*-based system for directed evolution of GPCRs in mammalian cells"

### Supporting Information Text

#### Tailored Library Design

For directed evolution of larger proteins such as GPCRs error-prone PCR is commonly used to introduce mutations. While this has proven to work for a number of receptors (1-5), some limitations of this technique must be considered. Unbiased randomization will inevitably lead to incorporation of unwanted mutations such as premature stop codons, the stochastic nature of error-prone PCR makes adjacent three-base changes extremely unlikely, limiting the accessible mutation through the biasing of the genetic code, and tuning the mutational load over the whole gene is difficult. Moreover, it has become increasingly clear that contacts within and between the transmembrane helices contribute most to receptor stability and function, and such residues can now be easily inferred from available protein structures or homology models. Therefore, a more directed design for GPCR libraries becomes possible which will improve the outcome of directed evolution.

To implement this, we applied a semi-rational approach for a tailored library design. For NTR1, the crystal structure of the thermostabilized variant NTR1-TM86V (PDB ID: 4BUO) was used to design the positions of mutations, but the only mutation permanently encoded in the library was R167$^{350}$L (see below). Residues that are facing into the receptor transmembrane bundle, and thus most likely contribute to allosteric interactions within the receptor core, were selected for randomization. Residues facing into the lipidic environment, or constituting the orthosteric binding pocket, or likely being involved in G protein interaction, were excluded. As a result, 94 positions were assigned for randomization.

Next, a GPCR-specific evolutionary substitution matrix was created which resembled commonly used matrices that are based on the frequencies of amino acid substitutions observed in aligned protein sequences (6-8). For this purpose, a multiple sequence alignment from 296 class A GPCRs (excluding olfactory receptors) was obtained from GPCRdb (see below), and the five most frequent amino acids at each position were determined to serve as library members for subsequent randomization (**Fig. S1**). Notably, the structural template NTR1-TM86V contains 11 stabilizing mutations. With the exception of R167$^{350}$L (c.f. main text), none of these mutations were permanently encoded into the library.

For PTH1R, a slightly modified strategy was applied (**Fig. S3**). Considering the complex multi-domain structure, inherent in class B GPCRs, randomization was restricted exclusively to the TMD, thereby omitting alterations to the ECD which primarily would affect full-length ligand binding. To identify residues for randomization, a homology model based on the structure of glucagon receptor (PDB ID: 4L6R) was used, since at the time, no structure for PTH1R had been available. Then, by applying the same algorithm as for NTR1, 99 positions were assigned for randomization. In contrast to the class A of GPCRs, the group of class B receptors is relatively small, and hence instead of an evolutionary model as for NTR1 we used a substitution matrix based on amino acid similarity (**Fig. S3B**). In addition, we assigned 19 positions that had been evolved in a previous directed evolution approach of PTH1R in yeast (2) (Klenk et al. unpublished). For those residues, the same randomization matrix was applied, yet specific mutations, which had been identified in the yeast selection but were not included in the substitution matrix, were added manually (e.g. M312K, K359N, Q440R; **Fig. S3C**).

To enable site- and sequence-specific randomization, cDNA libraries for NTR1 and PTH1R were created by Slonomics™ solid phase synthesis (9-11). Binomial distribution was set to 3-5 mutations per gene, requiring a mutagenesis rate of ~1% for each of the 5 substituents.

### Supplementary methods

#### Ligands

Human PTH(1–34) and PTH(3–34) were from Bachem. Neurotensin 8–13 [NT(8–13)] was from Anaspec. Human [Ac5c$^{1}$, Aib$^{3}$, Q$^{10}$, Har$^{11}$, A$^{12}$, W$^{14}$]PTH(1–14), [M-PTH(1–14)], was synthesized by Peptide Specialty Laboratories. M-PTH(1–14) was labeled at K$^{13}$ with HiLyte dye 647 [M-PTH(1–14)-HL647]. [Nle$^{8,18}$,Y$^{34}$,C$^{35}$]PTH(1–34) was labeled at C$^{35}$ with HiLyte dye 647 [PTH'(1–34)-HL647, the prime indicating the sequence changes compared to PTH(1–34)]. Neurotensin 8–13 was fluorescently labeled with HiLyte-647 [HL647-NT(8–13)] or HiLyte-488 [HL488-NT(8–13)] at the N-terminal amino group. All fluorescently labeled peptides were custom synthesized by Anaspec.

#### Cell Culture

HEK293T cells (Open Biosystems), A-431 and BSC-1 cells (ATCC) were cultivated in Dulbecco's modified medium supplemented with 10% (v/v) fetal calf serum. Cells were maintained at 37°C in a humidified atmosphere of 5% $CO_2$, 95% air. Transient transfection of HEK293T cells was performed with TransIT-293 (Mirus) reagent according to the manufacturer's protocol. CHO-S cells (Life Technologies) were maintained as shaking suspension culture using Power CHO 2CD medium (Lonza) supplemented with 8 mM L-glutamine, 0.1 mM hypoxanthine and 0.1 mM thymidine. Cells were seeded in DMEM with 10% fetal calf serum overnight to facilitate adhesion before transfection. Transient transfection was performed with Lipofectamine reagent (Invitrogen) according to the manufacturer's protocol.

#### NTR1 DNA Library Cloning

Acceptor constructs for rat NTR1, containing a cloning cassette with the *Mus musculus* IgG signal sequence aa 1-17, were constructed for both the mammalian expression plasmid (EFMOD, Vaccinex, 5' BssHII and 3' SalI cloning sites) and *Vaccinia* transfer plasmid (VHEH5, Vaccinex, 5' BssHII and 3' BsiWI cloning sites). The wild-type rat NTR1 gene (amino acids 43 – 424), along with NTR1 variants, L5X and TM86V, were amplified using PCR (iProof high-fidelity DNA polymerase, BioRad) and the following primers: NTRBSSHIIsense 5'-ttttt*GCGCGC*ACTCCACCTCGGAATCCGACACGG-3' and NTRaddSal1 5'-tttt*GTCGAC*TCAGTACAGGGTCTCCCGGGTG-3' (for EFMOD) or NTRAddBsiW1stop-5'-tttttCGTACGtTCAGTACAGGGTCTC- 3' (for VHEH5) by standard protocols. PCR products were subsequently cloned into the expression and transfer plasmids. DNA library construction was performed by PCR (Advantage2 polymerase, Clontech) of the mutant library 2218_-1_LIB_rNTR1 (43-424) using the same primers. The PCR product was resolved on 1% agarose/TBE gels. The 1177 bp band was gel-purified (Qiaquick, Qiagen), digested and ligated into the VHEH5 plasmid using NxGenT4 DNA ligase (Lucigen). High-efficiency transformation was done by electroporation of NEB10Beta *E. coli* cells (BioRad GenePulser, 1 mm cuvette, 2.0 kV, 200 Ω, 25 µF) to create a plasmid library with a diversity of ~1.4 × $10^{7}$.

**PTH1R DNA library DNA cloning**

Acceptor constructs for human PTH1R with a cloning cassette containing the PTH1R signal sequence aa 1–23 (including a naturally occurring BsiWI site) and a 3' SalI site were constructed for both mammalian expression (EFMOD, Vaccinex) and the inducible *Vaccinia* transfer plasmid (T7terVHE, Vaccinex). The full-length wild-type human PTH1R gene (1–593) were amplified using PCR (Q5 DNA polymerase, NEB) and cloned into expression and transfer plasmids (BsiWI/SalI). DNA library construction was performed by PCR (Q5 DNA Polymerase, NEB) of the linear DNA of mutant library SLN2248 with standard conditions and minimal cycling using the following primers: PTH1Rsignalsense 5'-CTCAGCTCCGCGTACGCGCTGGTG-3' and PTH1R AS 5'-TGTCCGTTCGGTCGACTCACATGACTGTCTCC-3'. The PCR product was resolved on 1% agarose/TBE gels. The 1734 bp band was gel purified (Qiaquick, Qiagen), digested with BsiWI/ SalI and ligated into T7TerVHE plasmid using NxGenT4 DNA ligase (Lucigen). High-efficiency transformation was described above to create a plasmid library with a diversity of ~$6.3 \times 10^6$.

**Generation of Vaccinia Virus Clones and Variant Libraries**

*Vaccinia* vector V7.5 virus (Vaccinex) was digested with Proteinase K (Thermo Fisher), and DNA was purified by phenol/chloroform extraction. V7.5 viral DNA was digested with restriction endonucleases ApaI (NEB) and NotI (NEB) and purified with Amicon ultra centrifugal columns (Millipore Sigma). BSC-1 cells were infected with helper fowlpox virus at a multiplicity of infection (MOI) of 1.5 plaque forming units (pfu) per cell and transfected with digested V7.5 vector DNA and each receptor library and the corresponding control plasmids. Infected/ transfected cells were incubated for 5 days, and *Vaccinia* virus was harvested by freeze-thawing the cells. Individual plaques for control clones were picked and amplified. Viral DNA was purified and amplified by PCR. Positive clones were confirmed by sequencing. Virus stock for the library was titered by plaque assay. Individual clones were randomly picked, and checked by PCR for recombination efficiency. The resulting *Vaccinia* libraries had over 95% positive recombinant efficiency harboring ~$1.3 \times 10^8$ and ~$1.1 \times 10^8$ unique recombinants for NTR1 and PTH1R, respectively.

***Vaccinia* Virus Infection and Fluorescent Ligand Binding**

A-431 cells were seeded the day before infection in DMEM + 10% (v/v) FBS and allowed to double overnight at 37°C, 5% $CO_2$. Cells were then infected overnight at an MOI of 1 pfu per cell with virus expressing NTR1 controls or the library. The appropriate pfu of virus was diluted into a minimal medium volume to cover the cell monolayer and incubated at 37°C, 5% $CO_2$ for 1-2 hours. Cells were then overlaid with sufficient media and allowed to incubate for 16 – 18 hours. Cells were harvested using Accutase™, pelleted and washed in FACS buffer [PBS, 1% (w/v) BSA] or Tris buffer [20 mM Tris-HCl (pH 7.4), 118 mM NaCl, 5.6 mM glucose, 1.2 mM $KH_2PO_4$, 1.2 mM $MgSO_4$, 4.7 mM KCl, 1.8 mM CaCl, 0.1% (w/v) BSA]. Cells were resuspended at $2 \times 10^6$ cells per ml in FACS buffer or Tris buffer and incubated with fluorescent ligand on ice for 1 – 2 hours. To confirm specificity, duplicate cell samples were also incubated with a 100-fold excess of unlabeled ligand. Cells were then washed in the appropriate buffer and fixed in 0.5% paraformaldehyde with propidium iodide for live/ dead cell discrimination before analysis on the flow cytometer.

**Fluorescence Activated Cell Sorting of Improved NTR1 Variants**

A-431 cells, infected with a library of NTR1 clones, were sorted for multiple iterations using 40 nM NT(8–13)-HL647. For the first round of sorting, $4 \times 10^7$ A-431 cells were infected with the NTR1 library at an MOI of 1 overnight at 37°C, 5% $CO_2$, as described above. The next day, the cells were harvested using Accutase™ and stained with 40 nM of the ligand in 1 ml total volume FACS buffer on ice for one hour with occasional, gently swirling. Cells were then washed twice with FACS buffer, resuspended at 2 × $10^7$ cells per ml and passed through a 40 µm filter before being sorted on the BD FACS Aria sorter. The top 0.3% fluorescent cells were collected (6,800 total), lysed by multiple freeze/thaw cycles and the virus was amplified in multiple flasks of BSC-1 cells for 2–3 days.

The amplified virus was harvested and titered before being used to infect A-431 cells again for a second round of sorting. Since the diversity of the pool from the first sort was only 6,800, 3 × $10^6$ A-431 cells were infected for the second sort. The top 0.06% events were collected and amplified as above. Enrichment in the sort was tested by small-scale infections and ligand staining throughout.

**Fluorescence Activated Cell Sorting of Improved PTH1R Variants**

A-431 cells infected with a library of PTH1R clones were sorted for multiple iterations using 120 nM M-PTH(1–14)-HL647 or 120 nM PTH'(1–34)-HL647. For the first round of sorting, 1.2 × $10^8$ A-431 cells were infected with the PTH1R T7-inducible library and the attenuated T7 promoter virus at an MOI of 1 overnight at 37°C, 5% $CO_2$, as described above. The next day, cells were harvested using Accutase™ and stained with 120 nM of ligand in 6 ml total volume on ice for one hour with occasional, gently swirling. Cells were then washed twice with FACS buffer, resuspended at 2 × $10^6$ cells per ml and passed through a 40 µm filter before being sorted on a BD FACS Aria sorter. The top 0.5% fluorescent cells were collected, lysed by multiple freeze/ thaw cycles, and the virus was amplified in multiple flasks of BSC-1 cells for 2 – 3 days.

The amplified virus was harvested and titered before being used to infect A-431 cells again for a second round of sorting. Each subsequent round of sorting was performed with 1.5 × $10^7$ A-431 cells and gating stringency was increased for each round. Sort enrichment was tested as small-scale infections and ligand staining throughout.

**Isolation of Receptor Variant DNA and Cloning into Mammalian Expression Vectors**

*Vaccinia* DNA was extracted from the sorted pools (DNA Blood mini, Qiagen). Pool variants were amplified from the pool DNA using PCR (Advantage2 polymerase, Clontech) by standard protocols with minimal cycling. For NTR1, primers NTRBSSHIIsense 5'-tttt*GCGCGC*ACTCCACCTCGGAATCCGACACGG-3' and NTRaddSal1 5'-tttt*GTCGAC*TCAGTACAGGGTCTCCCGGGTG-3' (EFMOD), and for PTHR1, signal sense 5'-CTCAGCTCCGCGTACGCGCTGGTG-3' and PTHR1AS 5'-CCCCCCTCGAGGTCGACTCACATGACTGTCTCCC-3'. PCR products were subsequently cloned into the mammalian expression vector EFMOD. Mini-libraries were prepared by picking 92 – 94 colonies and isolating plasmid DNA (Qiaprep 96 turbo, Qiagen). DNA sequences were analyzed by Sanger sequencing using 2 – 3 primers for full coverage.

**Preparation of mini-$G_s$ Protein**

Mini-$G_s$ 393 was essentially prepared as described before (12). Briefly, mini-$G_s$ was expressed in *E. coli* strain BL21(DE3) at 20°C. Cells were harvested 16–20 h post-induction by centrifugation, resuspended in lysis buffer [40 mM HEPES (pH 7.5), 150 mM NaCl, 5 mM imidazole, 10% (v/v) glycerol, 5 mM $MgCl_2$, 50 µM GDP, 1 mM DTT, 50 µg/ml DNAseI, 50 µg/ml lysozyme] and disrupted in a HPL6 cell lyser (Maximator GmbH) at 1700 bar. Lysates were clarified by centrifugation (20,000*g* for 45 min) and supernatants were loaded on Ni-NTA columns (Thermo Fisher Scientific).

Columns were washed with 10 CV wash buffer [20 mM HEPES (pH 7.5), 500 mM NaCl, 28 mM imidazole, 10% (v/v) glycerol, 1 mM $MgCl_2$, 50 µM GDP, 1 mM DTT], and bound proteins were eluted stepwise in 2 CV of elution buffer [20 mM HEPES (pH 7.5), 150 mM NaCl, 500 mM imidazole, 10% (v/v) glycerol, 1 mM $MgCl_2$, 50 µM GDP, 0.5 mM DTT]. Imidazole was removed on PD-10 desalting columns (Cytiva), proteins were concentrated to 25 mg/ml and snap-frozen in freezing buffer [25 mM HEPES (pH 7.5), 150 mM NaCl, 15% (v/v) glycerol, 5 mM $MgCl_2$, 10 µM GDP, 0.25 mM DTT].

**Expression Analysis**

Receptor variants were transiently transfected in HEK293T cells. 48 hrs after transfection, cells were detached with Accutase and incubated with 20 nM HL488-NT(8–13) or 100 nM PTH'(1–34)-HL647 in PBS supplemented with 0.2% BSA for 2 – 4 h on ice. Nonspecific binding was determined in the presence of a 100-fold excess of unlabeled peptide. Cells were then washed three times with ice-cold PBS, and fluorescence intensities were determined on a FACSCanto II flow cytometer (BD Biosciences).

**Ligand Binding Assays**

Ligand binding experiments were performed on whole cells or on cell membranes obtained from transiently transfected HEK293T cells, using in both cases an HTRF binding assay as described before (2, 13). All receptor variants were subcloned into a mammalian expression vector containing an N-terminal SNAP-tag (Cisbio). For NTR1 constructs, the SNAP tag was fused to residue 43 of the receptor. For PTH1R, the SNAP tag was either fused to residue 29 or to residue 171, thus eliminating the ECD. HEK293T cells were transiently transfected with receptor constructs and were seeded at 20,000 cells per well in poly-L-lysine-coated 384-well plates (Greiner) for whole-cell binding assays or at $5 \times 10^6$ cells in 10 cm Petri dishes for membrane preparation. 48 h after transfection, cells were incubated with 50 nM SNAP-Lumi4-Tb (Cisbio) in ligand binding buffer [20 mM HEPES pH 7.5, 100 mM NaCl, 3 mM $MgCl_2$ and 0.02% (w/v) BSA] for 2 h at 37 °C. Cells were washed four times with assay buffer and used directly for whole cell ligand binding experiments, or crude cell membrane extracts were prepared as described before. Cells or 0.2–1 µg membranes per well were then incubated for 4 h on ice to measure ligand binding, containing fluorescently labeled tracer peptide together with a concentration range of unlabeled competitor peptide. For NTR1, 2 nM of HL488-NT(8–13) was used as a tracer peptide. For PTH1R, 50 nM of M-PTH(1–14)-HL647 or 20 nM of PTH'(1–34)-HL647 were used. Fluorescence intensities were measured on a Spark fluorescence plate reader (Tecan) with an excitation wavelength of 340 nm and emission wavelengths of 620 nm, 520 nm and 665 nM for $Tb^{3+}$, HiLyte Fluor 488 and HiLyte Fluor 647, respectively. The ratio of FRET-donor and -acceptor fluorescence intensities was calculated. Total binding was obtained in the absence of competitor, and nonspecific binding was determined in the presence of a 100-fold excess of unlabeled peptide. Data were normalized to the specific binding for each individual experiment and were analyzed by global fitting to a one-site heterologous competition equation.

**Signaling Assays**

Signaling experiments were performed on whole cells with transiently transfected HEK293T cells as described before (2, 13). Twenty-four hours after transfection, cells were washed with PBS, detached with cell dissociation buffer (Gibco) and washed again in PBS. Cells were resuspended in assay buffer [10 mM Hepes (pH 7.4), 146 mM NaCl, 1 mM $CaCl_2$, 0.5 mM $MgCl_2$, 4.2 mM KCl,

5.5 mM glucose, 50 mM LiCl, 1 mM 3-isobutyl-1-methylxanthin). cAMP and IP1 accumulation assays were performed on white low-volume 384-well plates (Greiner) using the cAMP Tb kit and the IP-One Tb kit (both from CisBio), respectively, according to the manufacturer's protocol. For cAMP accumulation, 5,000 cells were incubated with agonist at the indicated concentrations for 30 min at RT. To determine basal receptor signaling, cells were incubated in IBMX-containing assay buffer for 30 min in the absence of ligand. For IP1 accumulation, 20,000 cells were incubated with agonist at the indicated concentrations for 2 h at 37°C. Fluorescence intensities were measured on a Spark fluorescence plate reader (Tecan). To generate concentration-response curves, data were fitted to a three-parameter logistic equation.

**Thermostability Measurements**

Stability of evolved receptor variants was measured in membrane fractions of transiently transfected HEK293T cells by determining the residual receptor-bound ligand after a heat challenge of the membranes. In the case of PTH1R, the ECD (residues 1–170) was removed from the expression constructs to restrict stability measurements to the TMD. For NTR1, cells were left unmodified, whereas for PTH1R, cells were labeled with 50 nM SNAP-Lumi-4Tb before membranes were prepared as described above. Membranes were then incubated for 2–4 h on ice in ligand-binding buffer containing 20 nM of [3,11-tyrosyl-3,5-$^{3}$H(N)]-neurotensin (Perkin Elmer) and 500 nM of M-PTH(1–14)-HL647 for NTR1 and for PTH1R, respectively. Where indicated, 25 µM of mini-$G_s$ protein were added to the membrane fractions prior to ligand addition. Thereafter, 0.5 µg of membranes were distributed per well of a 96-well plate and heated to a specific temperature in a PCR thermocycler for 20 min. NTR1-containing membranes were then immobilized on glass fiber filters (Millipore), washed four times with binding buffer, and the residual activity of the radio-ligand was measured on a MicroBeta Plus 1450 liquid scintillation counter (Perkin Elmer). For PTH1R, residual ligand binding was determined by HTRF as described above. Data were analyzed by nonlinear regression fitting.

**Data Quantification, Statistical Analysis and Visualization**

Flow cytometry data were analyzed in FlowJo software 10 (BD Biosciences). All other statistical analysis and curve fitting was performed in Prism 6.07 (GraphPad). Details of each analysis are outlined in the experimental methods section, figures, tables and figure legends of the specific experiment. Sequence alignments and snake plots were obtained from the GPCRdb (14). Sequence frequencies were visualized with WebLogo (15). Statistically significant differences were determined by one-way ANOVA and Bonferroni multi-comparison.

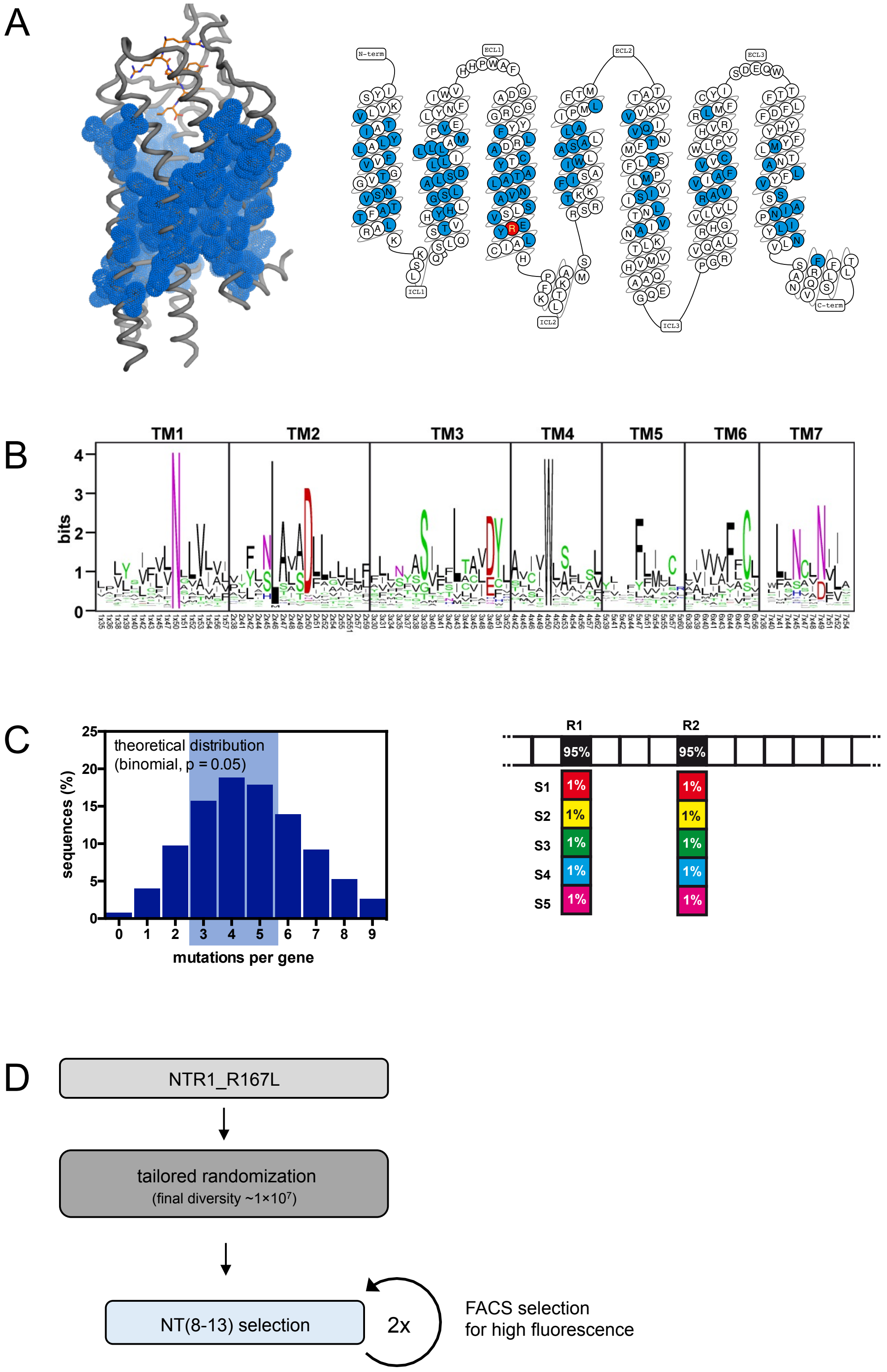


**Fig. S1.** Library design and selection strategy for NTR1. (*A*) Based on the crystal structure of rNTR1 (PDB ID: 4BUO), 94 residues were selected for randomization. R167$^{3.50}$L was included as a fixed mutation to the library (red circle). (*B*) Substituents were selected based on an evolutionary substitution matrix. For this purpose, a multiple sequence alignment from 296 class A GPCRs (excluding olfactory receptors) was obtained, and the five most frequent amino acids at each position were determined to serve as library members for subsequent randomization. (*C*) A mutation frequency was chosen to yield an average number of 3–5 mutations per gene. For this purpose, per residue each of the 5 substituents was introduced with 1% frequency. (*D*) Selection scheme for NTR1. After incorporating the library into the *Vaccinia* vector, A-431 cells were infected and 2 consecutive selection rounds with 40 nM HL647-NT(8–13) were performed.

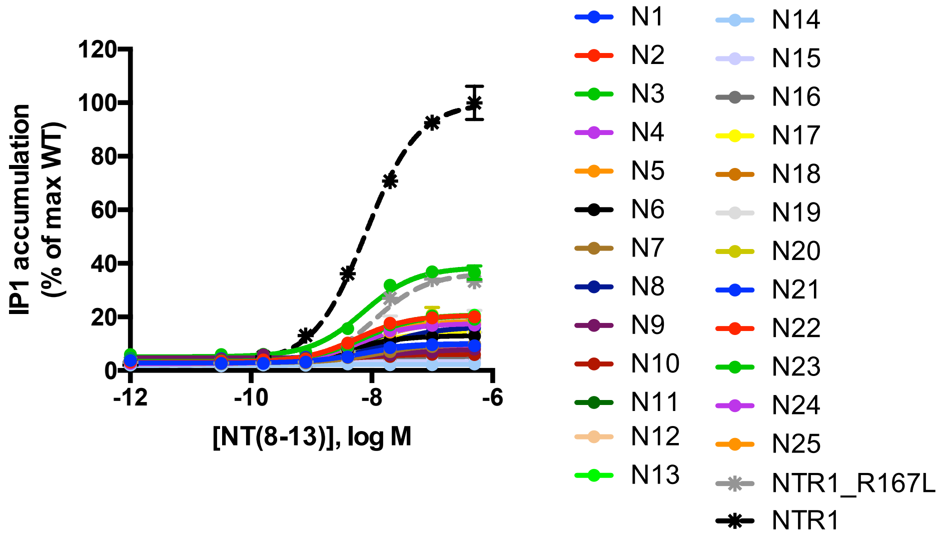


**Fig. S2.** Signaling activity of NTR1 mutants. For each receptor variant, $G_q$-mediated IP1 accumulation was measured in HEK293T cells after stimulation with NT(8–13). Data were normalized to IP1 levels of NTR1 wild type at 500 nM NT(8–13) and are shown as mean values (± s.e.m.) of 2 independent experiments each performed in duplicates.

A

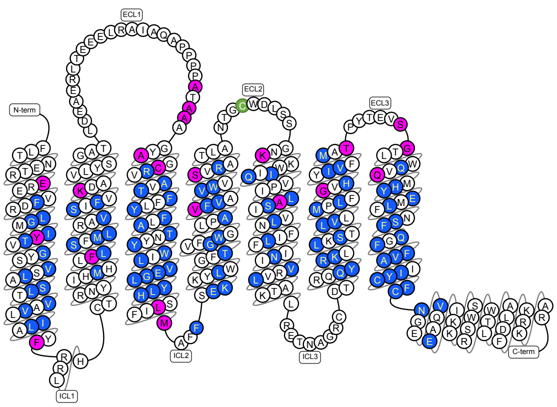


B

| | | | | | | | | | | | | | | | | | | |
|---|---|---|---|---|---|---|---|---|---|---|---|---|---|---|---|---|---|---|
| **WT** | **E** | **K** | **R** | **H** | **T** | **S** | **N** | **Q** | **G** | **A** | **C** | **V** | **I** | **L** | **M** | **F** | **Y** | **W** |
| **S1** | D | H | H | F | A | A | C | C | A | C | A | A | A | A | A | H | C | F |
| **S2** | S | Q | K | K | C | C | D | E | C | I | N | C | F | F | I | I | F | H |
| **S3** | T | R | Q | R | N | N | Q | N | S | L | S | I | L | I | L | L | H | I |
| **S4** | Q | I | F | W | S | T | S | S | V | S | T | L | M | M | Q | W | L | L |
| **S5** | A | M | L | Y | V | V | T | T | L | V | V | T | V | V | V | Y | W | Y |

C

| | | | | | | | | | | | | | | | | | | | |
|---|---|---|---|---|---|---|---|---|---|---|---|---|---|---|---|---|---|---|---|
| | TM1 | TM1 | TM2 | ECL1 | ECL1 | ECL1 | ECL1 | TM3 | TM3 | ICL2 | TM4 | TM4 | ECL1 | ECL2 | TM5 | TM6 | TM6 | ECL3 | ECL3 | TM7 |
| WN | 1.32 | 1.43 | 2.67 | | | | | 3.29 | 3.57 | | 4.55 | 4.62 | | | 5.45 | 6.50 | 6.59 | | | 7.38 |
| AA | 180 | 191 | 240 | 272 | 274 | 275 | 279 | 281 | 309 | 312 | 334 | 341 | 351 | 359 | 369 | 418 | 427 | 433 | 434 | 440 |
| **WT** | E | Y | K | A | A | A | A | C | L | M | V | S | C | K | A | G | T | S | G | Q |
| **S1** | D | C | H | C | C | C | C | A | A | A | A | A | A | E | C | A | A | A | A | C |
| **S2** | S | F | Q | I | I | I | I | S | F | I | C | C | S | H | I | C | C | C | C | N |
| **S3** | T | H | R | L | L | L | L | T | H | K | I | I | T | N | L | D | I | L | S | R |
| **S4** | Q | L | I | S | S | S | S | V | I | L | L | N | V | Q | S | S | S | N | V | S |
| **S5** | A | W | M | V | V | V | V | Y | V | Q | T | T | Y | R | V | V | V | T | W | T |

D

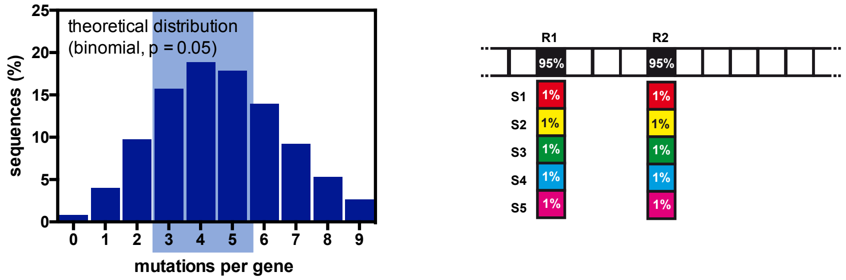


E

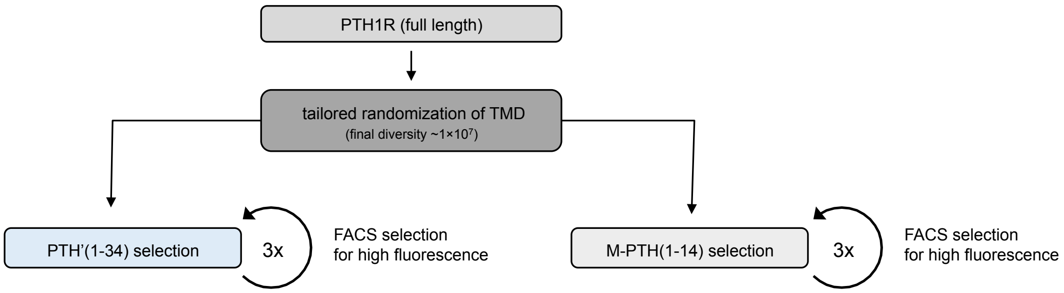


**Fig. S3.** Library design and selection strategy for PTH1R. (*A*) A homology model for PTH1R based on the crystal structure of glucagon receptor (PDB ID: 4L6R) was generated, and 99 residues within the TMD (residues 171-480) were selected for randomization (blue). Additionally, 19 residues identified in a previous evolution campaign in yeast (2) (Klenk et al. unpublished) (magenta) as well as the conserved Cys351 in ECL2 (green) were randomized. (*B*) Substitution matrix for 18 amino acids. Asp and Pro were not among the WT residues assigned for randomization and thus are not contained in the matrix. Each of the five alternative codons for randomization (S1 – S5) is based on amino acid similarity to the wild-type (WT) amino acid. (C) Yeast-derived residues and Cys351 were randomized following the same scheme as in (B) with the exception that stabilizing amino acids (black outline) were included in the substitution matrix. AA, residue number; WN, residue number according to Wootten (16). (*D*) The mutation frequency was chosen to yield an average distribution of 3 – 5 mutations per gene. For this purpose, at each residue position, each of the five alternative codons (S1 – S5) was incorporated with 1% frequency. (*E*) Selection scheme for PTH1R. After transferring the library into the *Vaccinia* vector, A-431 cells were infected and three consecutive selection rounds were performed either with 120 nM PTH'(1–34)-HL647 or with 120 nM M-PTH(1–14)-HL647.

A

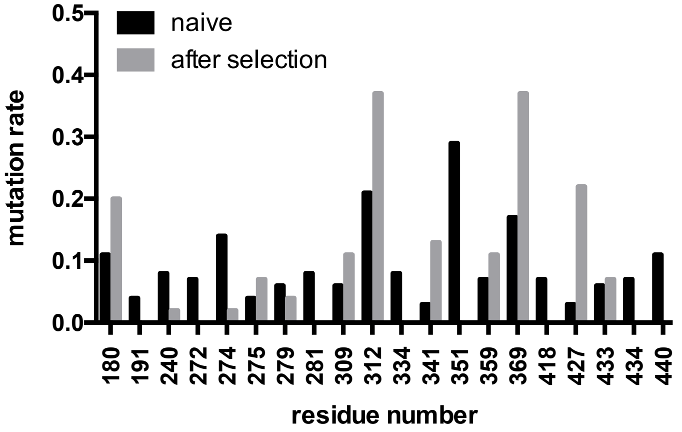


B

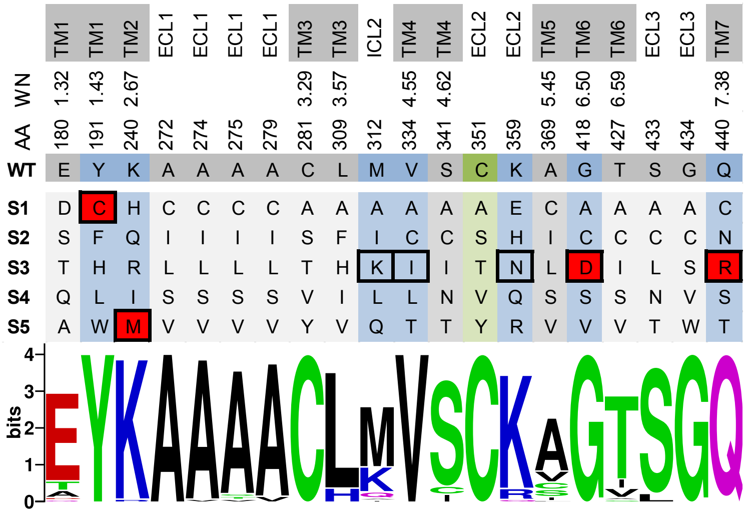


**Fig. S4.** Mutations that disrupt PTH1R signaling are deselected. (*A*) Mutation rate of the naive library and after three selection rounds. 96 sequences of each pool were analyzed. Shown are 19 positions, which had been derived from a previous yeast evolution campaign to stabilize PTH1R (2) (Klenk et al. unpublished), and position 351, which is required for disulfide bond formation between ECL2 and ECL3 in wild-type PTH1R (2) (c.f. **Fig. S3A**). (*B*) Amino acid distribution of 92 clones after three selection rounds. The wild-type sequence (WT) and the initial randomization scheme (S1 to S5) (c.f. **Fig. S3B-D**) are shown in the top panel. Stability-conferring positions identified in the yeast evolution campaign are shaded in blue and the respective stabilizing mutation is marked by a black outline. Stabilizing mutations Y191$^{1.43}$C, K240$^{2.67}$M, G418$^{6.50}$D and Q440$^{7.38}$R that disrupted receptor signaling (2) are shaded in red. C351 required for disulfide bond formation between ECL2 and ECL3 is shaded in green. The sequence logo shows the amino acid distribution of 92 clones after the selection at the positions indicated, indicating that receptor-inactivating mutations have been deselected. AA, residue number; WN, residue number according to Wootten (16)

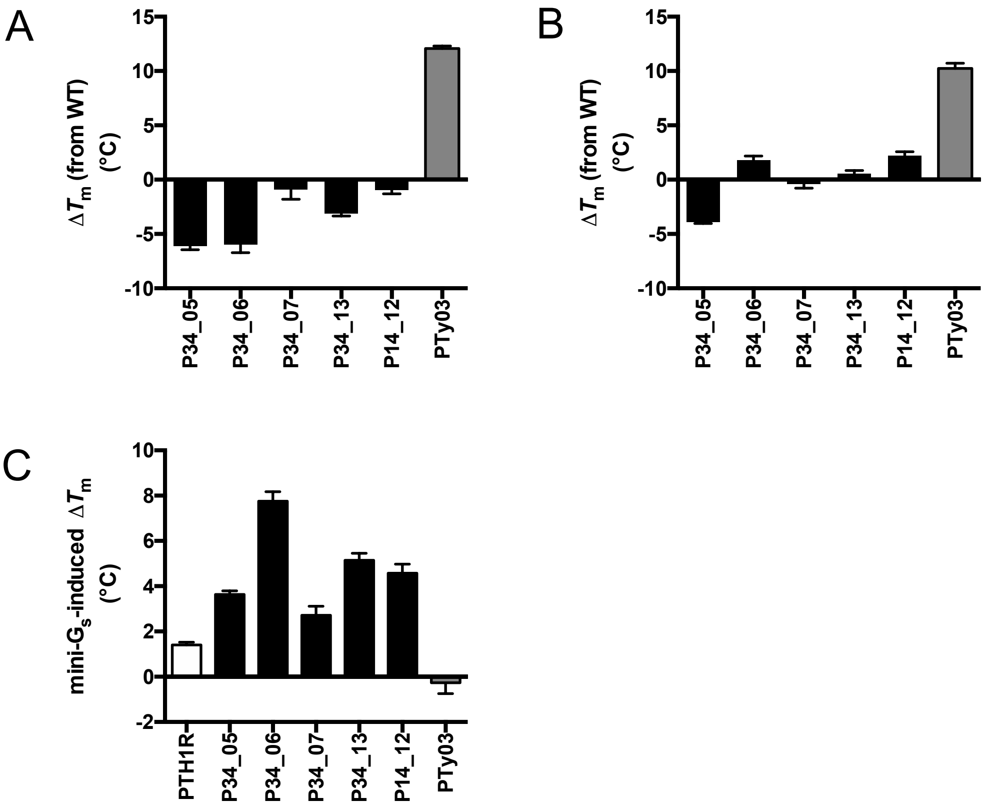


**Fig. S5.** Thermostability of evolved PTH1R variants is G protein-dependent. Thermostability of PTH1R variants was assessed in membrane preparations in the absence (*A*) or presence (*B*) of 12.5 µM mini-$G_s$. Data are given as the change in $T_m$ from wild-type PTH1R. (*C*) Change in $T_m$ induced by the presence of G protein. Data are shown as the change in $T_m$ from each variant in absence of mini-$G_s$. The thermostabilized, signaling-inactive variant PTy03 (2) was included as control (grey). Data repesent mean values ± s.e.m. of 4 independent experiments (**Table S5**).

**Table S1 |** Pharmacological data of evolved NTR1 variants.

| | expression (fold of WT) | NT(8–13) binding $pIC_{50}$ (log M) | IP1 signaling $pIC_{50}$ (log M) | $E_{max}$ (% WT) |
|---|---|---|---|---|
| N1 | 47.8 ± 10.7 (2) | 8.35 ± 0.03 (3) | 8.15 ± 0.13 (2) | 7.0 ± 0.5 (2) |
| N2 | 54.2 ± 4.3 (2) | 8.34 ± 0.04 (3) | 8.25 ± 0.08 (2) | 17.6 ± 0.7 (2) |
| N3 | 43.7 ± 2.2 (2) | 8.51 ± 0.10 (3) | 8.14 ± 0.07 (2) | 17.2 ± 0.6 (2) |
| N4 | 51.1 ± 2.3 (2) | 8.42 ± 0.04 (3) | 8.24 ± 0.05 (2) | 15.0 ± 0.4 (2) |
| N5 | 45.6 ± 1.5 (2) | 8.43 ± 0.03 (3) | 8.19 ± 0.09 (2) | 15.9 ± 0.7 (2) |
| N6 | 49.5 ± 3.7 (2) | 8.58 ± 0.06 (3) | 8.34 ± 0.08 (2) | 9.8 ± 0.4 (2) |
| N7 | 74.4 ± 7.1 (2) | 8.11 ± 0.05 (3) | 7.87 ± 0.09 (2) | 7.1 ± 0.3 (2) |
| N8 | 76.4 ± 0.6 (2) | 8.00 ± 0.05 (3) | 7.83 ± 0.06 (2) | 13.4 ± 0.5 (2) |
| N9 | 36.1 ± 2.0 (2) | 8.27 ± 0.05 (3) | 7.50 ± 0.40 (2) | 3.2 ± 0.8 (2) |
| N10 | 40.2 ± 2.0 (2) | 7.93 ± 0.08 (3) | 8.31 ± 0.23 (2) | 2.7 ± 0.3 (2) |
| N11 | 49.2 ± 8.6 (2) | 8.35 ± 0.07 (3) | 8.11 ± 0.06 (2) | 15.7 ± 0.5 (2) |
| N12 | 36.0 ± 1.8 (2) | 8.16 ± 0.08 (3) | 8.13 ± 0.08 (2) | 15.0 ± 0.6 (2) |
| N13 | 44.4 ± 5.0 (2) | 8.37 ± 0.10 (3) | 8.20 ± 0.07 (2) | 17.7 ± 0.6 (2) |
| N14 | 82.5 ± 9.9 (3) | 8.07 ± 0.03 (3) | n.a. | 0.4 ± 0.2 (2) |
| N15 | 28.8 ± 8.3 (2) | 8.20 ± 0.02 (3) | 8.71 ± 0.34 (2) | 1.4 ± 0.2 (2) |
| N16 | 30.8 ± 1.9 (2) | 8.36 ± 0.07 (3) | 8.18 ± 0.36 (2) | 1.7 ± 0.3 (2) |
| N17 | 33.7 ± 6.8 (2) | 8.25 ± 0.06 (3) | 8.22 ± 0.08 (2) | 11.3 ± 0.5 (2) |
| N18 | 48.3 ± 12.9 (2) | 8.17 ± 0.08 (3) | 8.09 ± 0.09 (2) | 15.7 ± 0.7 (2) |
| N19 | 27.9 ± 2.7 (2) | 8.32 ± 0.05 (3) | 8.22 ± 0.18 (2) | 17.0 ± 1.5 (2) |
| N20 | 25.3 ± 0.3 (2) | 8.35 ± 0.02 (3) | 8.17 ± 0.13 (2) | 17.6 ± 1.1 (2) |
| N21 | 40.9 ± 1.7 (2) | 8.49 ± 0.03 (3) | 8.18 ± 0.18 (2) | 5.7 ± 0.5 (2) |
| N22 | 27.6 ± 0.1 (2) | 8.39 ± 0.07 (3) | 8.26 ± 0.09 (2) | 15.9 ± 0.7 (2) |
| N23 | 35.5 ± 4.1 (2) | 8.21 ± 0.06 (3) | 8.14 ± 0.08 (2) | 33.4 ± 1.4 (2) |
| N24 | 41.0 ± 1.5 (2) | 7.79 ± 0.09 (3) | 8.16 ± 0.17 (2) | 6.2 ± 0.5 (2) |
| N25 | 42.2 ± 1.0 (2) | 7.98 ± 0.02 (3) | 8.06 ± 0.09 (2) | 5.0 ± 0.2 (2) |
| NTR1 | 1.0 ± 0.0 (3) | 7.13 ± 0.07 (3) | 7.95 ± 0.07 (2) | 98.9 ± 1.8 (2) |
| NTR1_R167L | 2.5 ± 0.8 (3) | 7.57 ± 0.07 (3) | 7.95 ± 0.07 (2) | 34.4 ± 1.3 (2) |
| NTR1-TM86V | 47.2 ± 6.5 (2) | n.d. | n.d. | n.d. |
| NTR1-L5X | 33.4 ± 1.0 (2) | n.d. | n.d. | n.d. |

Expression levels were determined by flow cytometry using 20 nM HL488-NT(8–13). All data are represented as mean values ± s.e.m.. The number of experiments is given in parentheses. n.d., not determined; n.a. not applicable

**Table S2 |** Thermostability of evolved NTR1 variants

| | $T_m$ (°C) |
|---|---|
| N8 | 60.0 ± 0.7 (4) |
| N12 | 58.1 ± 0.6 (4) |
| N13 | 56.4 ± 0.6 (4) |
| N14 | 62.4 ± 0.6 (4) |
| N15 | 59.8 ± 0.5 (4) |
| N21 | 57.7 ± 0.4 (4) |
| N23 | 57.9 ± 0.4 (4) |
| NTR1_R167L | 51.4 ± 0.6 (4) |
| NTR1 | 52.5 ± 0.6 (4) |

Thermostability data were obtained by measuring loss of ligand binding as a function of temperature in membrane fractions All data are represented as mean values ± s.e.m.. The number of independent experiments is given in parentheses.

**Table S3 |** Expression and ligand binding of evolved PTH1R variants.

| | **expression (fold of WT)** | **M-PTH(1–14) binding $pIC_{50}$ (log M)** | **PTH(1–34) binding $pIC_{50}$ (log M)** |
|---|---|---|---|
| P14_01 | 6.3 ± 1.6 (2) | 7.86 ± 0.70 (5) | 7.86 ± 0.09 (2) |
| P14_02 | 3.6 ± 0.1 (2) | 7.19 ± 0.09 (3) | 7.83 ± 0.08 (2) |
| P14_03 | 8.9 ± 0.4 (2) | 7.83 ± 0.14 (3) | 8.07 ± 0.11 (2) |
| P14_04 | 1.8 ± 0.1 (2) | 7.26 ± 0.02 (3) | 7.73 ± 0.01 (2) |
| P14_05 | 5.2 ± 1.7 (2) | 7.14 ± 0.02 (3) | 7.82 ± 0.20 (4) |
| P14_06 | 2.5 ± 0.7 (2) | 7.28 ± 0.01 (3) | 7.32 ± 0.03 (2) |
| P14_07 | 1.2 ± 0.0 (2) | 7.77 ± 0.06 (3) | 8.40 ± 0.18 (2) |
| P14_08 | 2.3 ± 0.4 (2) | 7.58 ± 0.02 (3) | 8.32 ± 0.17 (2) |
| P14_09 | 1.7 ± 0.6 (2) | 7.50 ± 0.11 (3) | 7.82 ± 0.01 (2) |
| P14_10 | 4.1 ± 0.5 (2) | 7.79 ± 0.09 (4) | 8.05 ± 0.11 (2) |
| P14_11 | 3.3 ± 0.6 (2) | 7.06 ± 0.15 (5) | 7.98 ± 0.03 (2) |
| P14_12 | 3.6 ± 0.5 (2) | 7.25 ± 0.50 (4) | 8.09 ± 0.06 (2) |
| P14_13 | 4.4 ± 0.4 (2) | 6.83 ± 0.49 (3) | 8.31 ± 0.08 (2) |
| P14_14 | 1.8 ± 0.3 (2) | 7.85 ± 0.20 (3) | 8.32 ± 0.07 (2) |
| P14_15 | 3.6 ± 0.2 (2) | 7.67 ± 0.12 (3) | 8.06 ± 0.01 (2) |
| P14_16 | 2.4 ± 0.4 (2) | 7.85 ± 0.33 (3) | 8.50 ± 0.24 (2) |
| P34_01 | 8.0 ± 2.2 (2) | 7.46 ± 0.04 (3) | 7.97 ± 0.02 (2) |
| P34_02 | 9.5 ± 2.0 (2) | 7.59 ± 0.06 (6) | 7.95 ± 0.04 (2) |
| P34_03 | 7.1 ± 1.1 (2) | 7.02 ± 0.06 (4) | 7.69 ± 0.04 (2) |
| P34_04 | 8.5 ± 1.6 (2) | 7.70 ± 0.06 (3) | 8.04 ± 0.16 (2) |
| P34_05 | 5.3 ± 1.4 (2) | 7.31 ± 0.05 (3) | 7.89 ± 0.12 (2) |
| P34_06 | 7.5 ± 0.1 (2) | 8.55 ± 1.12 (5) | 8.07 ± 0.10 (2) |
| P34_07 | 12.5 ± 2.1 (2) | 7.74 ± 0.07 (3) | 7.99 ± 0.10 (2) |
| P34_08 | 2.7 ± 0.3 (2) | 7.62 ± 0.13 (3) | 7.90 ± 0.01 (2) |
| P34_09 | 5.9 ± 0.9 (2) | 7.88 ± 0.07 (3) | 7.95 ± 0.06 (2) |
| P34_10 | 4.6 ± 1.1 (2) | 7.23 ± 0.03 (3) | 8.09 ± 0.16 (2) |
| P34_11 | 3.5 ± 0.7 (2) | 7.01 ± 0.04 (3) | 7.83 ± 0.10 (2) |
| P34_12 | 4.6 ± 1.7 (2) | 7.58 ± 0.08 (3) | 8.07 ± 0.26 (2) |
| P34_13 | 6.2 ± 1.2 (2) | 7.34 ± 0.07 (3) | 7.87 ± 0.10 (2) |
| P34_14 | 1.4 ± 0.1 (2) | 7.46 ± 0.07 (5) | 8.09 ± 0.03 (2) |
| P34_15 | 1.2 ± 0.3 (2) | 7.11 ± 0.05 (4) | 7.98 ± 0.06 (3) |
| P34_16 | 1.5 ± 0.3 (2) | 7.58 ± 0.10 (5) | 7.84 ± 0.03 (2) |
| P34_17 | 1.5 ± 0.0 (2) | 7.86 ± 0.15 (3) | 8.28 ± 0.03 (3) |
| P34_18 | 5.6 ± 0.7 (2) | 7.66 ± 0.03 (4) | 8.56 ± 0.00 (2) |
| P34_19 | 7.4 ± 1.2 (2) | 7.63 ± 0.12 (3) | 8.40 ± 0.00 (2) |
| P34_20 | 3.8 ± 1.2 (2) | 7.62 ± 0.06 (5) | 8.33 ± 0.02 (2) |
| P34_21 | 2.3 ± 0.1 (2) | 6.99 ± 0.03 (3) | 7.85 ± 0.04 (2) |

| | | | |
|---|---|---|---|
| P34_22 | 1.6 ± 0.1 (2) | 7.58 ± 0.06 (3) | 8.44 ± 0.15 (2) |
| P34_23 | 2.2 ± 0.5 (2) | 7.51 ± 0.06 (3) | 8.17 ± 0.01 (2) |
| P34_24 | 0.9 ± 0.0 (2) | 6.25 ± 0.10 (3) | 8.27 ± 0.08 (2) |
| P34_25 | 1.8 ± 0.1 (2) | 8.28 ± 0.49 (3) | 8.60 ± 0.20 (2) |
| P34_26 | 1.9 ± 0.2 (2) | 7.17 ± 0.60 (3) | 8.40 ± 0.05 (2) |
| P34_27 | 5.7 ± 1.1 (2) | 4.86 ± 1.31 (5) | 8.11 ± 0.12 (2) |
| PTH1R | 1.0 ± 0.0 (2) | 6.23 ± 0.08 (7) | 7.80 ± 0.09 (8) |

Expression levels were determined by flow cytometry analysis using PTH'(1–34)-HL647. HTRF-ligand binding assays were performed on whole cells. M-PTH(1–14) binding was determined in constructs only containing the TMD of receptor whereas PTH(1–34) binding was obtained in full-length receptor constructs. All data are represented as mean values ± s.e.m.. The number of independent experiments is given in parentheses.

**Table S4 |** cAMP accumulation of evolved PTH1R variants

| | **p$IC_{50}$ (log M)** | **$E_{max}$ (fold of WT)** |
|---|---|---|
| P14_01 | 9.37 ± 0.05 (4) | 1.19 ± 0.23 (4) |
| P14_02 | 10.42 ± 0.36 (3) | 1.06 ± 0.17 (3) |
| P14_03 | 9.89 ± 0.18 (4) | 1.24 ± 0.22 (4) |
| P14_04 | 10.31 ± 0.01 (3) | 1.86 ± 0.50 (3) |
| P14_05 | 10.75 ± 0.21 (5) | 0.79 ± 0.23 (5) |
| P14_06 | 10.48 ± 0.48 (3) | 0.87 ± 0.33 (3) |
| P14_07 | 10.51 ± 0.10 (3) | 0.95 ± 0.28 (3) |
| P14_08 | 10.22 ± 0.09 (3) | 1.41 ± 0.58 (3) |
| P14_09 | 10.01 ± 0.08 (2) | 1.85 ± 0.65 (2) |
| P14_10 | 9.84 ± 0.27 (1) | 0.57 ± 0.12 (3) |
| P14_11 | 10.32 ± 0.04 (2) | 1.10 ± 0.25 (2) |
| P14_12 | 10.42 ± 0.50 (2) | 0.57 ± 0.14 (2) |
| P14_13 | 9.79 ± 0.44 (2) | 1.00 ± 0.14 (2) |
| P14_14 | 10.34 ± 0.35 (2) | 0.72 ± 0.16 (2) |
| P14_15 | 9.22 ± 0.31 (2) | 1.10 ± 0.21 (2) |
| P14_16 | 9.79 ± 0.18 (2) | 1.06 ± 0.22 (2) |
| P34_01 | 10.30 ± 0.43 (3) | 1.12 ± 0.28 (3) |
| P34_02 | 10.13 ± 0.06 (6) | 1.39 ± 0.31 (6) |
| P34_03 | 10.25 ± 0.43 (3) | 1.74 ± 0.35 (3) |
| P34_04 | 9.93 ± 0.17 (3) | 1.61 ± 0.28 (3) |
| P34_05 | 10.13 ± 0.46 (3) | 1.55 ± 0.28 (3) |
| P34_06 | 10.31 ± 0.32 (3) | 1.55 ± 0.25 (3) |
| P34_07 | 9.22 ± 0.08 (3) | 1.49 ± 0.42 (3) |
| P34_08 | 10.60 ± 0.47 (4) | 1.58 ± 0.92 (4) |
| P34_09 | 10.53 ± 0.23 (2) | 1.64 ± 0.19 (2) |
| P34_10 | 10.82 ± 0.48 (2) | 1.75 ± 0.01 (2) |
| P34_11 | 10.17 ± 0.34 (2) | 1.81 ± 0.46 (2) |
| P34_12 | 9.79 ± 0.41 (2) | 1.48 ± 0.19 (2) |
| P34_13 | 10.93 ± 0.40 (2) | 2.17 ± 0.66 (2) |
| P34_14 | 10.21 ± 0.10 (2) | 1.43 ± 0.29 (2) |
| P34_15 | 10.04 ± 0.24 (5) | 1.24 ± 0.28 (5) |
| P34_16 | 10.95 ± 0.21 (1) | 1.11 ± 0.05 (3) |
| P34_17 | 8.88 ± 1.53 (1) | 2.16 ± 0.32 (3) |
| P34_18 | 10.48 ± 0.33 (4) | 0.82 ± 0.06 (4) |
| P34_19 | 10.43 ± 0.17 (2) | 0.61 ± 0.08 (2) |
| P34_20 | 10.24 ± 0.39 (2) | 0.83 ± 0.13 (2) |
| P34_21 | 9.73 ± 0.11 (2) | 1.24 ± 0.06 (2) |
| P34_22 | 9.72 ± 0.01 (2) | 0.32 ± 0.13 (2) |

| | | |
|---|---|---|
| P34_23 | 10.50 ± 0.19 (2) | 0.46 ± 0.10 (2) |
| P34_24 | 10.06 ± 0.41 (5) | 1.00 ± 0.04 (5) |
| P34_25 | 10.02 ± 0.30 (2) | 0.90 ± 0.34 (2) |
| P34_26 | 10.10 ± 0.37 (2) | 0.86 ± 0.32 (2) |
| P34_27 | 9.90 ± 0.05 (1) | 0.81 ± 0.11 (3) |
| PTH1R | 10.49 ± 0.13 (5) | 1.00 ± 0.00 (5) |

cAMP accumulation was measured in transiently transfected HEK293T cells after stimulation with 1 μM PTH(1–34). All data are represented as mean values ± s.e.m.. The number of independent experiments is given in parentheses.

**Table S5 |** Thermostability of evolved PTH1R variants

| | **$T_m$ (°C)** | |
|---|---|---|
| | **- mini-$G_s$** | **+ mini-$G_s$** |
| P34_05 | 44.7 ± 0.2 (4) | 48.3 ± 0.2 (4) |
| P34_06 | 46.2 ± 0.5 (4) | 53.8 ± 0.4 (4) |
| P34_07 | 48.9 ± 0.8 (4) | 51.6 ± 0.4 (4) |
| P34_13 | 47.5 ± 0.3 (4) | 52.7 ± 0.3 (4) |
| P14_12 | 49.7 ± 0.4 (4) | 54.5 ± 0.4 (4) |
| PTH1R | 50.6 ± 1.0 (4) | 52.0 ± 0.1 (4) |
| PTy03 | 62.7 ± 0.3 (4) | 62.7 ± 0.5 (4) |

Thermostability data were obtained by measuring loss of ligand binding as a function of temperature in membrane fractions in the absence or presence of 12.5 μM mini-$G_s$. All data are represented as mean values ± s.e.m.. The number of independent experiments is given in parentheses.

### SI References


1. C. A. Sarkar *et al.*, Directed evolution of a G protein-coupled receptor for expression, stability, and binding selectivity. *Proc. Natl. Acad. Sci. U. S. A.* **105**, 14808–14813 (2008).

2. J. Ehrenmann *et al.*, High-resolution crystal structure of parathyroid hormone 1 receptor in complex with a peptide agonist. *Nat. Struct. Mol. Biol.* **25**, 1086–1092 (2018).

3. I. Dodevski, A. Plückthun, Evolution of Three Human GPCRs for Higher Expression and Stability. *J. Mol. Biol.* **408**, 599–615 (2011).

4. M. Schütz *et al.*, Directed evolution of G protein-coupled receptors in yeast for higher functional production in eukaryotic expression hosts. *Sci. Rep.* **6**, 21508 (2016).

5. Y. Waltenspühl, J. R. Jeliazkov, L. Kummer, A. Plückthun, Directed evolution for high functional production and stability of a challenging G protein-coupled receptor. *Sci. Rep.* **11**, 8630 (2021).

6. S. Henikoff, J. G. Henikoff, Amino acid substitution matrices from protein blocks. *Proc. Natl. Acad. Sci. U. S. A.* **89**, 10915–10919 (1992).

7. M. O. Dayhoff, R. M. Schwartz, B. C. Orcutt, "A Model of Evolutionary Change in Proteins" in *Atlas of Protein Sequence and Structure*, M. O. Dayhoff, Ed. (National Biomedical Research Foundation Silver Spring MD, 1978), pp. 345–352.

8. S. Rios *et al.*, GPCRtm: An amino acid substitution matrix for the transmembrane region of class A G Protein-Coupled Receptors. *BMC Bioinformatics* **16**, 206 (2015).

9. J. Van den Brulle *et al.*, A novel solid phase technology for high-throughput gene synthesis. *BioTechniques* **45**, 340–343 (2008).

10. W. Zhai *et al.*, Synthetic antibodies designed on natural sequence landscapes. *J. Mol. Biol.* **412**, 55–71 (2011).

11. K. M. Schlinkmann *et al.*, Maximizing detergent stability and functional expression of a GPCR by exhaustive recombination and evolution. *J. Mol. Biol.* **422**, 414–428 (2012).

12. B. Carpenter, C. G. Tate, Engineering a minimal G protein to facilitate crystallisation of G protein-coupled receptors in their active conformation. *Protein Eng. Des. Sel.* **29**, 583–594 (2016).

13. M. Deluigi *et al.*, Complexes of the neurotensin receptor 1 with small-molecule ligands reveal structural determinants of full, partial, and inverse agonism. *Sci. Adv.* **7** (2021).

14. A. J. Kooistra *et al.*, GPCRdb in 2021: integrating GPCR sequence, structure and function. *Nucleic Acids Res.* **49**, D335–D343 (2021).

15. G. E. Crooks, G. Hon, J.-M. Chandonia, S. E. Brenner, WebLogo: a sequence logo generator. *Genome Res.* **14**, 1188–1190 (2004).

16. D. Wootten, J. Simms, L. J. Miller, A. Christopoulos, P. M. Sexton, Polar transmembrane interactions drive formation of ligand-specific and signal pathway-biased family B G protein-coupled receptor conformations. *Proc. Natl. Acad. Sci. U. S. A.* **110**, 5211–5216 (2013).
